# Enhanced Cerebral Blood Volume under Normobaric Hyperoxia in the J20-hAPP Mouse Model of Alzheimer’s Disease

**DOI:** 10.1101/848713

**Authors:** Osman Shabir, Paul Sharp, Monica A Rebollar, Luke Boorman, Clare Howarth, Stephen B Wharton, Sheila E Francis, Jason Berwick

**Author notes:** Corresponding Author University of Sheffield, Western Bank, Sheffield, S10 2TN (United Kingdom).

## Abstract

Early impairments to neurovascular coupling have been proposed to be a key pathogenic factor in the onset and progression of Alzheimer’s disease (AD). Studies have shown impaired neurovascular function in several mouse models of AD, including the J20-hAPP mouse. In this study, we aimed to investigate early neurovascular changes using wild-type (WT) controls and J20-hAPP mice at 6-9 months of age, by measuring cerebral haemodynamics and neural activity to physiological sensory stimulations. A thinned cranial window was prepared to allow access to cortical vasculature and imaged using 2D-optical imaging spectroscopy (2D-OIS). After chronic imaging sessions where the skull was intact, a terminal acute imaging session was performed where an electrode was inserted into the brain to record simultaneous neural activity. We found that cerebral haemodynamic changes were significantly enhanced in J20-hAPP mice compared with controls in response to physiological stimulations, potentially due to the significantly higher neural activity (hyperexcitability) seen in the J20-hAPP mice. Thus, neurovascular coupling remained preserved under a chronic imaging preparation. Further, under hyperoxia, the baseline blood volume and saturation of all vascular compartments in the brains of J20-hAPP mice were substantially enhanced compared to WT controls, but this effect disappeared under normoxic conditions. This study highlights novel findings not previously seen in the J20-hAPP mouse model, and may point towards a potential therapeutic strategy by driving an increased baseline blood flow to the brain, thereby potentially enhancing the clearance of beta-amyloid.

## Introduction

Alzheimer’s disease (AD) is the most prevalent form of dementia worldwide and is characterised by a progressive decline in cognition. AD is pathologically characterised by the presence of extracellular amyloid beta (Aβ) plaques and intracellular neurofibrillary tangles composed of hyperphosphorylated-tau, which are associated with the progressive neurodegeneration and synaptic dysfunction seen in AD^1^.

At present there are limited disease modifying or curative treatments for AD and studying disease mechanisms in human subjects is difficult. Therefore, pre-clinical models of AD; mainly mouse models, have been generated to study AD mechanisms in vivo. Whilst numerous mouse models of AD exist, they do not fully recapitulate the human disease in its entirety^2,3^. However, these mouse models can effectively model specific aspects of AD pathology, such as amyloid plaque deposition and toxicity.

The J20-hAPP mouse model of AD over-expresses human amyloid precursor protein (hAPP) with the Swedish (K670N and M671L) and the Indiana (V7171F) familial mutations^4^. These mice produce more Aβ1-42 and plaques begin to readily form in the hippocampus from around 5-6 months of age^4,5^. The J20-hAPP mouse model displays significant neuroinflammation characterised by gliosis of both astrocytes and microglia^5^. They also display significant long-term memory impairment^5^.

The brain is extremely metabolically demanding, and the neurophysiological process of neurovascular coupling ensures that neurons receive an efficient and adequate blood supply to match the metabolic demands that neurons exert. The neurovascular degeneration hypothesis; as proposed by Zlokovic^6,7^, suggests that neurovascular breakdown is an important step in the pathogenesis of cerebrovascular and neurodegenerative disease, especially in AD. Evidence suggests that vascular dysregulation is the earliest feature of late-onset AD, preceding Aβ deposition, metabolism and structural deficits^8^. Therefore, studying neurovascular coupling and neurovascular degeneration is important to identify early biomarkers or treatment strategies.

Using a chronic mouse preparation, previous research from our laboratory found no significant neurovascular deficits in the J20-hAPP mouse between 9-12m age^9^, despite neuroinflammation and memory deficits^5^. This is contrary to what other laboratories have shown with the J20-mouse at the same age^10–12^. Such deficits have only been reported in acute experimental preparation sessions where the measurement of neurovascular function is performed on the same day as surgery, and not in chronic sessions where the effects of surgery have been mitigated. Based on these observations, we hypothesised that neurovascular function will not be altered in 6-9m old J20-AD mice using a chronic imaging preparation. The aim of the study therefore was to investigate neurovascular function at an earlier stage (between 6-9m) in the J20-hAPP mouse model to investigate whether there were neuronal or vascular abnormalities at an earlier disease stage when amyloid-plaques start to form, or whether they would remain intact as seen in 9-12m old J20-AD mice.

## Results

### Enhanced Blood Volume (HbT) Responses in J20-hAPP Mice

Imaging of the cortex through a thinned cranial window using 2D-OIS allows estimation of cortical haemodynamics in terms of relative changes of HbT (total haemoglobin, blood volume), HbR (deoxyhaemoglobin) & HbO (oxyhaemoglobin) concentration (Figure 1). Mechanical whisker stimulations at 5Hz evoke a haemodynamic response within the branches of the middle cerebral artery (MCA) and the immediately surrounding regions from which an active region of interest (ROI) can be determined (Figure 1B). From the ROI, an average time-series of the haemodynamics can be produced showing percentage changes of HbT, HbR & HbO over time; before, during and post-stimulation (Figure 1D).

**Figure 1:**
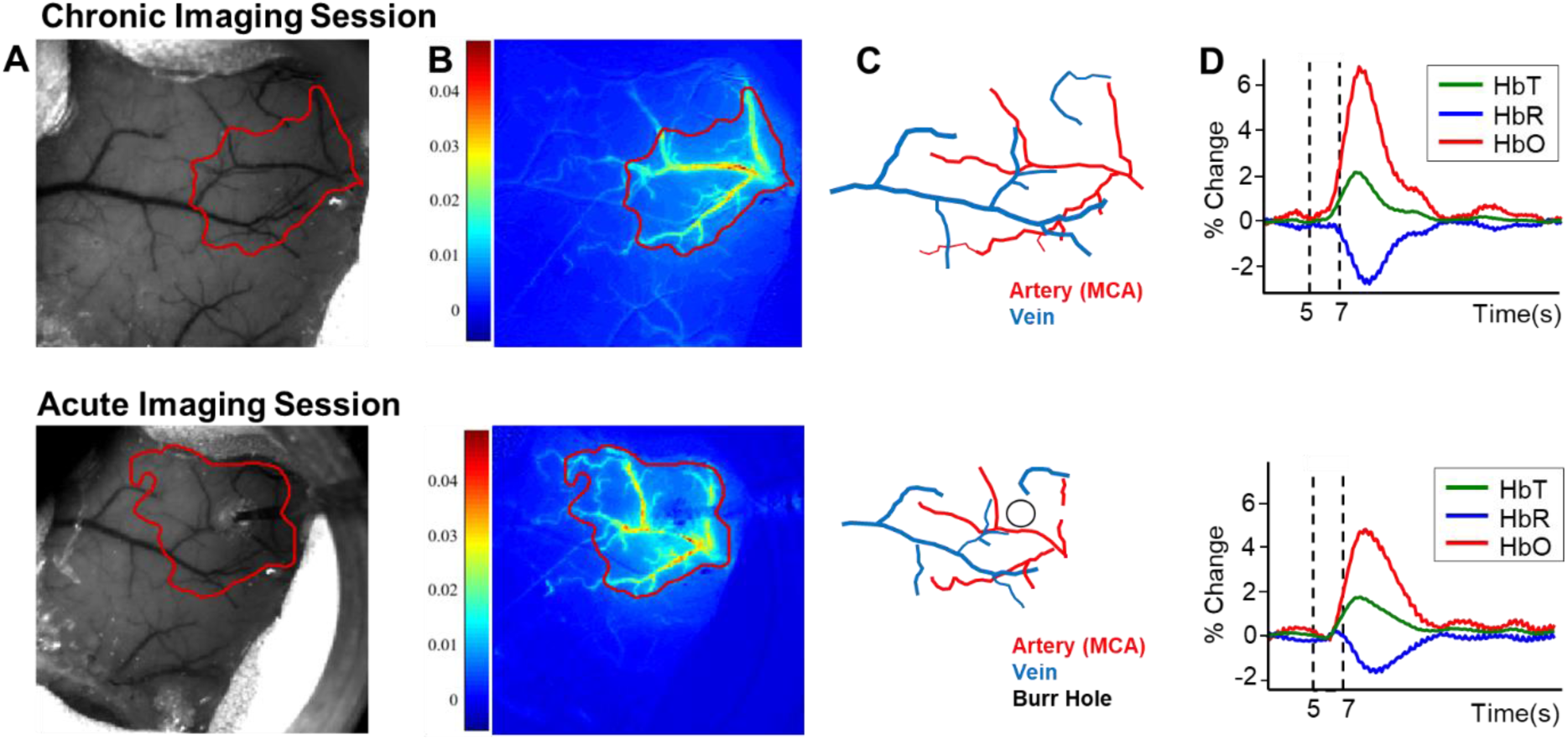
Representative 2D-OIS haemodynamic data from a WT mouse to a 2s-mechanical whisker stimulation. Top: chronic imaging sessions. Bottom: Acute imaging sessions with electrode. A) Raw grayscale image of thinned cranial window overlaid with the active region of interest (ROI; red) as defined from the spatial map of fractional changes in HbT (B). C) Vessel map showing arteries and veins within the window and active ROI. D) Time-series profiles of the haemodynamic data showing increases to HbT & HbO and a decrease (washout) in HbR. Dotted lines represent start and end time of whisker stimulation.

A week after chronic imaging sessions (in which the skull remains intact), a small burr-hole was drilled through the skull in the centre of the barrel cortex; determined from 2D-OIS data on a previous chronic imaging session, and a micro-electrode was inserted through the brain as a terminal experiment (Figure 1, bottom row). It is important to note that in acute imaging sessions the haemodynamic responses are typically smaller (Figure 1D).

Using the chronic thinned cranial window preparation (as shown in Figure 1-top) and imaging through an intact skull 2 weeks after surgery, the first question addressed was whether stimulation-evoked haemodynamic responses were different in J20-hAPP mice compared to WT controls. We found that stimulation-evoked HbT (blood volume) responses were significantly higher (on average by a +2% change in absolute values) in J20-AD mice compared to WT controls under both 100% O_2_ (hyperoxia) and 21% O_2_ (normoxia) conditions (2-way repeated-measures ANOVA of peak response: F=11.6, p=0.001) (Figure 2). HbO peak responses were also significantly increased (2-way repeated measures ANOVA: F=8.42, p=0.005). The washout of HbR was always significantly smaller in both WT and J20-AD mice in normoxic conditions (2-way repeated measures ANOVA: F=5.96, p=0.01), though not significantly more impaired in J20-AD mice compared to WTs (F=0.95, p=0.34). Irrespective of gas condition, HbT and HbO were always substantially higher in J20-hAPP mice compared to WTs in all experiments (Figure 2).

**Figure 2:**
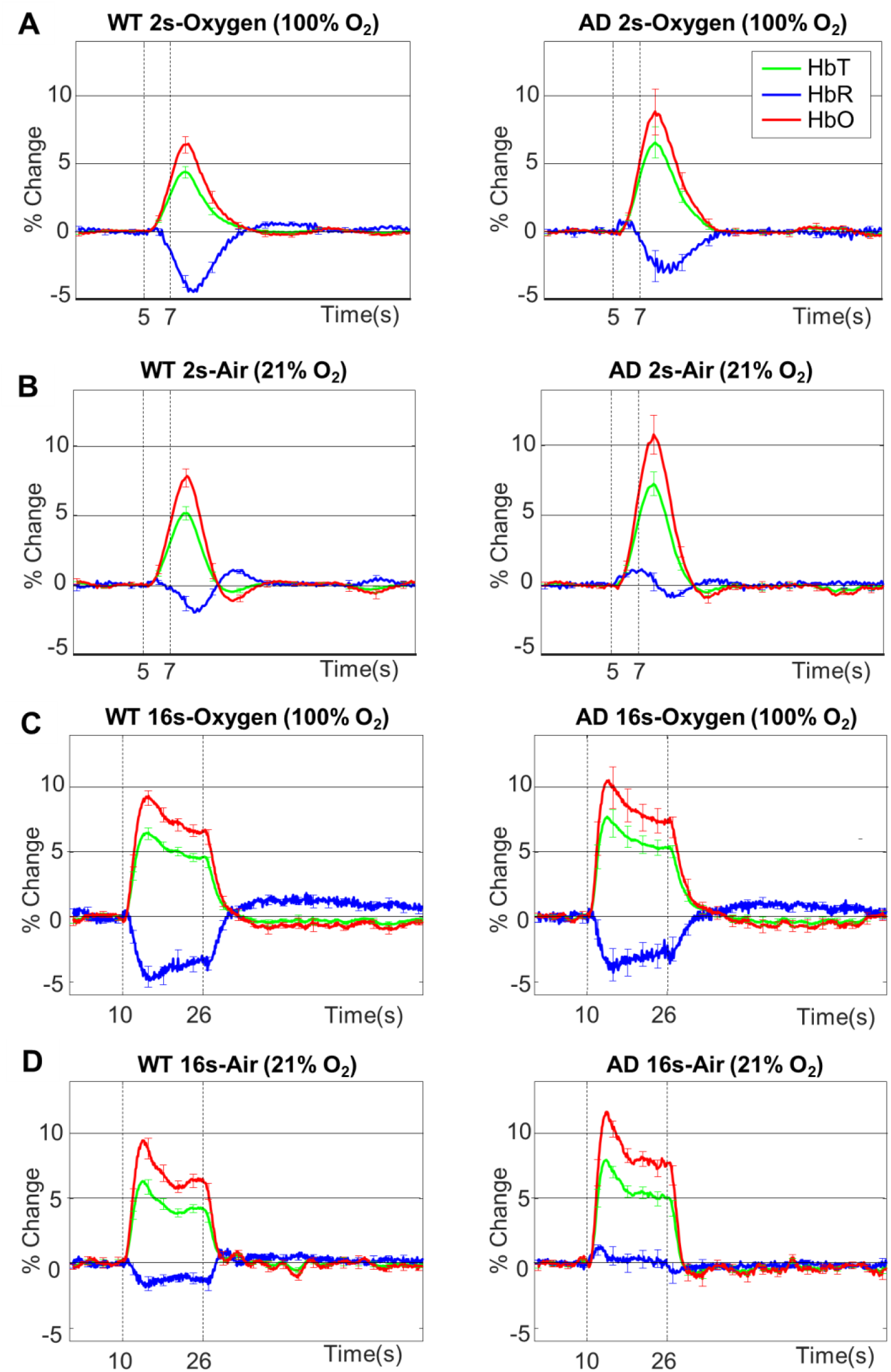
Mean stimulation-evoked haemodynamic responses in WT and J20-AD mice. WT (n=8) [left column] and J20-AD (n=9) [right column] with graphs showing mean % changes in the peak values of HbT, HbR & HbO ±SEM. **A)** 2s stimulation in 100% oxygen: WT-HbT 4.48±0.4, AD-HbT 6.7±1.09, WT-HbO 6.64±0.58, AD-HbO 9.06±1.57, WT-HbR −3.82±1.12, AD-HbR −3.99±1.09. **B)** 2s stimulation in 21% oxygen: WT-HbT 5.3±0.47, AD-HbT 7.38±0.8, WT-HbO 8.04±0.6, AD-HbO 11.04±1.25, WT-HbR −3.19±0.34, AD-HbR −1.62±0.4. **C)** 16s stimulation in 100% oxygen: WT-HbT 6.62±0.4, AD-HbT 8.09±1.04, WT-HbO 9.44±0.52, AD-HbO 11.04±1.55, WT-HbR −4.92±0.65, AD-HbR −4.15±1.15. **D)** 16s stimulation in 21% oxygen WT-HbT 6.44±0.49, AD-HbT 8.36±0.68, WT-HbO 9.64±0.68, AD-HbO 12.22±0.99, WT-HbR −2.06±0.42, AD-HbR −1.18±0.48. Dotted vertical lines indicate start and end time of whisker stimulation.

As part of the experimental paradigm, a test of global vascular reactivity was performed at the end of the experiment using 10% hypercapnia (Figure S1). We found no significant differences between WT and J20-AD mice (HbT: 2-tailed unpaired t-test p=0.53) suggesting that vessels in WT and J20-AD mice are able to dilate maximally to the same extent.

In 6 WT and 5 J20-hAPP mice, acute imaging sessions were performed where an electrode was inserted into the brain one week after chronic imaging sessions. In these sessions, there were no significant differences in the haemodynamic responses between WT and J20-hAPP mice: HbT (F=3.09, p=0.087), HbO (F=1.42, p=0.24) & HbR (F=3.73, p=0.06) (Figure S2).

### Neural Activity is Substantially Higher in J20-hAPP Mice

Next, we investigated whether the J20-hAPP mice had altered neural activity early in the disease course. We inserted a multichannel microelectrode into the centre of the barrel cortex (Figure 1-bottom row) and recorded neural data simultaneously with 2D-OIS. We found that stimulation-evoked multi-unit activity (MUA) was significantly higher in J20-hAPP mice compared to WT mice across all stimulation durations and gas conditions (2-way repeated measures ANOVA: F=8.82, p=0.005) (Figure 3). The enhanced neural activity was indicative of neural hyperexcitability in J20-AD mice. MUA responses were averaged from channels 3-8 (Figure 3A) which were at the depth of the somatosensory cortex.

**Figure 3.**
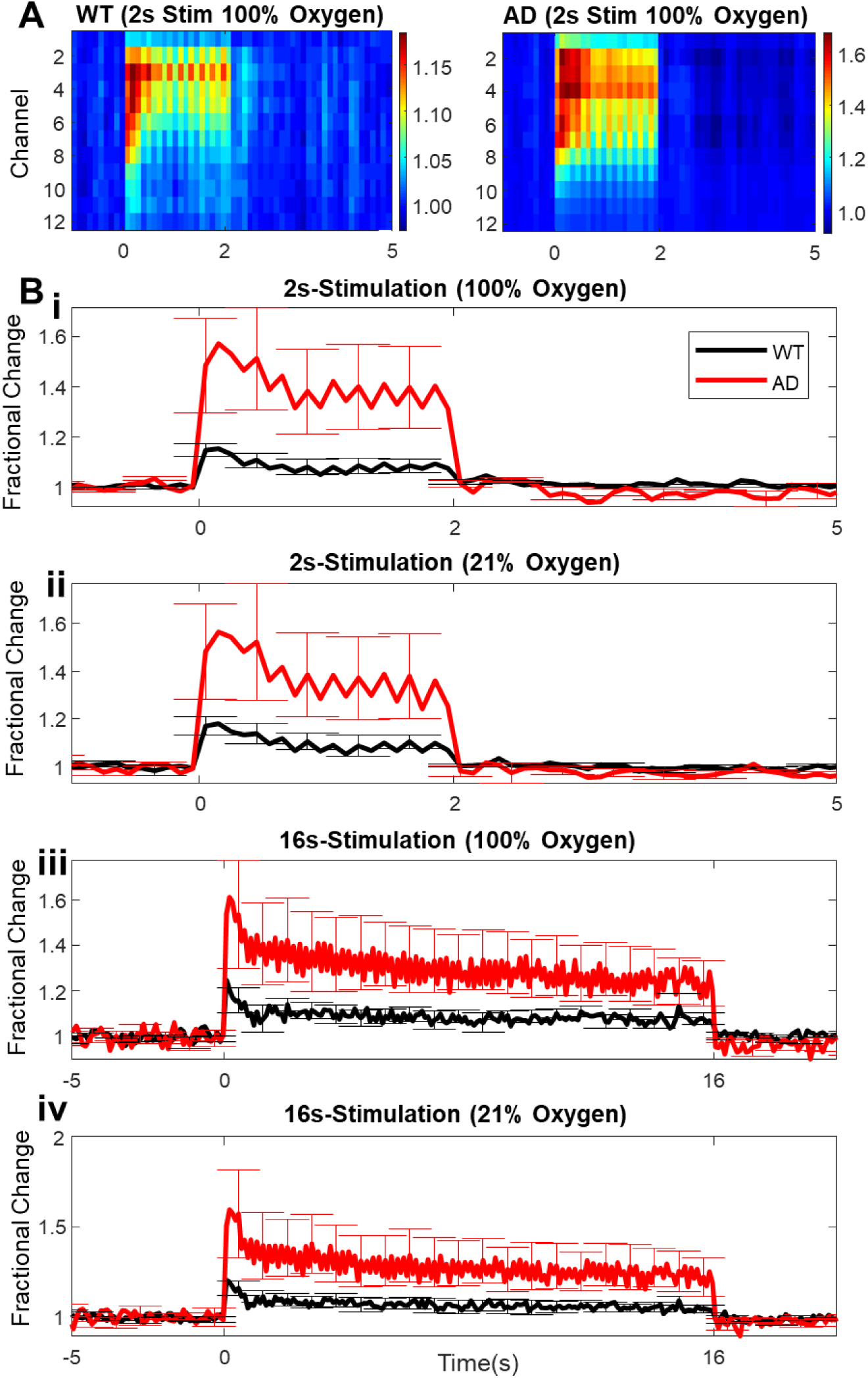
Multi-Unit Activity (MUA) Data. **A)** MUA activity (fractional changes of number of spikes/100ms) along depth of the microelectrode and time (stimulation periods) for 2s-stimulation in 100% oxygen. Cortical regions are at the depth of channels 3-8 (biggest responses in MUA). **B)** Time-series of mean total responses of MUA (fractional changes in spikes/100ms) ±SEM. **i)** 2s-stimulation in 100% oxygen: WT-MUA 547.52±98.7 spikes/100ms, AD-MUA 2260.78±710.8 spikes/100ms. **ii)** 2s-stimulation in 21% oxygen: WT-MUA 676.1±143.35 spikes/100ms, AD-MUA 2318.16±932.9 spikes/100ms. **iii)** 16s-stimulation in 100% oxygen: WT-MUA 4722.73±914.8 spikes/100ms, AD-MUA 14,158.4±4753.2 spikes/100ms. **iv)** 16s-stimulation in 21% oxygen: WT-MUA 3974.8±796.8 spikes/100ms, AD-MUA 13,592.8±5184.6 spikes/100ms. WT n=6, J20-hAPP n=5.

### Baseline Blood Volume is Substantially Higher in J20-AD Mice under Hyperoxia

As part of the experimental protocol, baseline haemodynamics during transition experiments (from 100% oxygen to 21% oxygen/air and vice versa) were recorded and quantified. We found that the baseline blood volume (HbT) was significantly enhanced in J20-AD mice under hyperoxic conditions (2-tailed unpaired t-test p=0.0002), but not in WT mice (Figure 4B). Upon transition from 100% oxygen to air (21% oxygen), there was a substantial decrease in HbT in J20-AD mice (−6.2% change on average) characterised by the enhanced vasoconstriction of all vascular compartments (Figure 4A/C).

**Figure 4.**
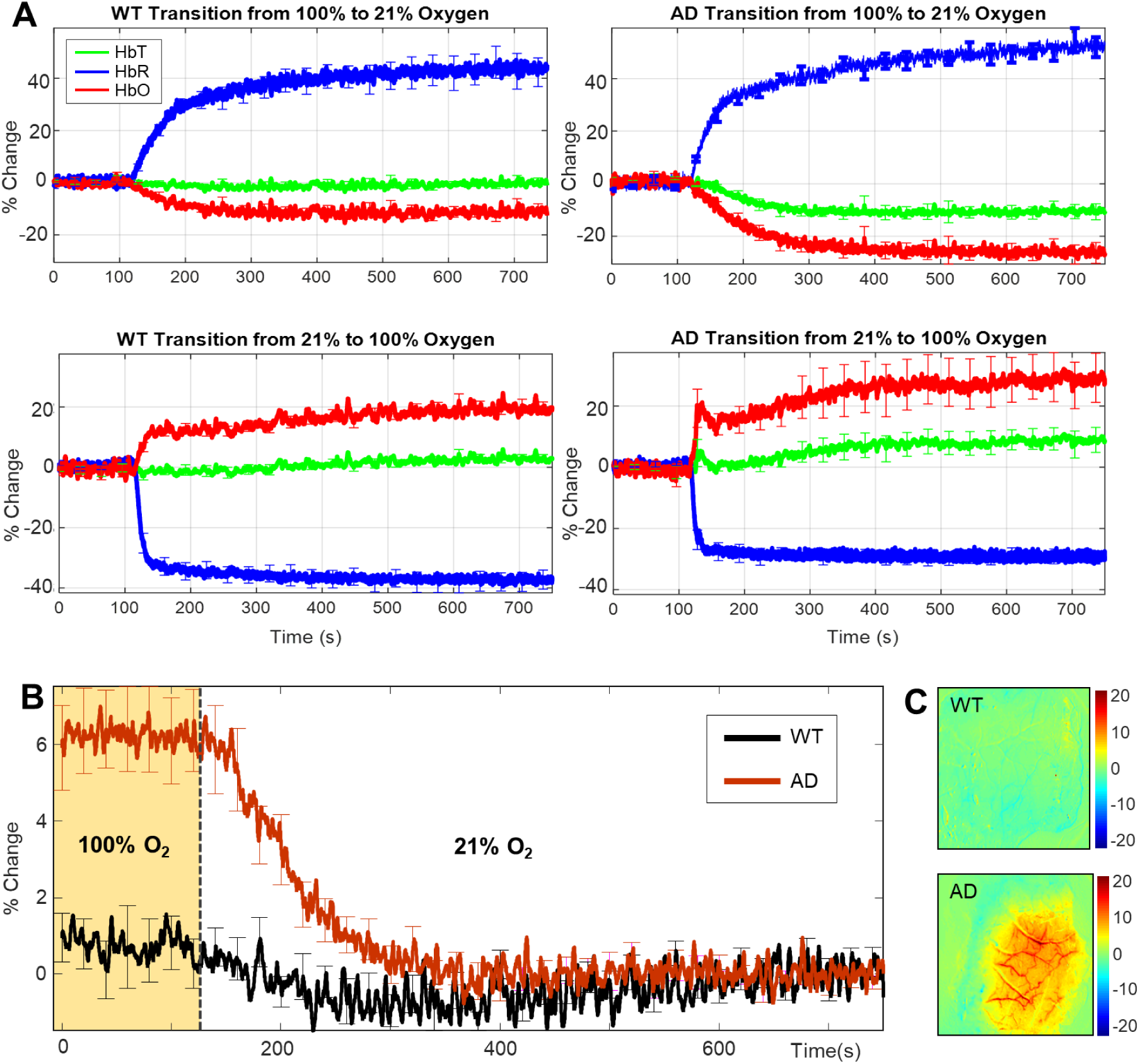
Average gas transitions between hyperoxic and normoxic conditions in WT and J20-AD mice. **A)** Time-series of gas transition experiments: top; transition from 100% oxygen to 21% oxygen & bottom; transition from 21% oxygen to 100% oxygen. **B)** Time-series of HbT at baseline at 100% oxygen and during gas transition to 21% oxygen with % change in HbT: WT-HbT 0.5%±1.8%STD, AD-HbT 6.2%%±3%STD. *Error bars ±SEM*. **(C)** Representative spatial map showing the differences in HbT across all vascular compartments in a J20-hAPP mouse (bottom), but not WT mouse (top) under normobaric hyperoxia.

### Amyloid-Plaques Begin to Form in the Hippocampus of 9m J20-AD Mice

Amyloid plaques are a key pathological hallmark of AD and begin to form in the hippocampus in the J20-hAPP AD mouse at around 6m of age^5^. The progression of plaque pathology and subsequent inflammatory changes have been well characterised in the J20-AD mouse^5^. We confirmed that in our cohort, by the age of 9m, the hippocampus has numerous medium sized plaques (Figure 5B) with fewer and smaller diffuse ones forming within the deeper cortex, but none within the upper layers of the cortex.

**Figure 5.**
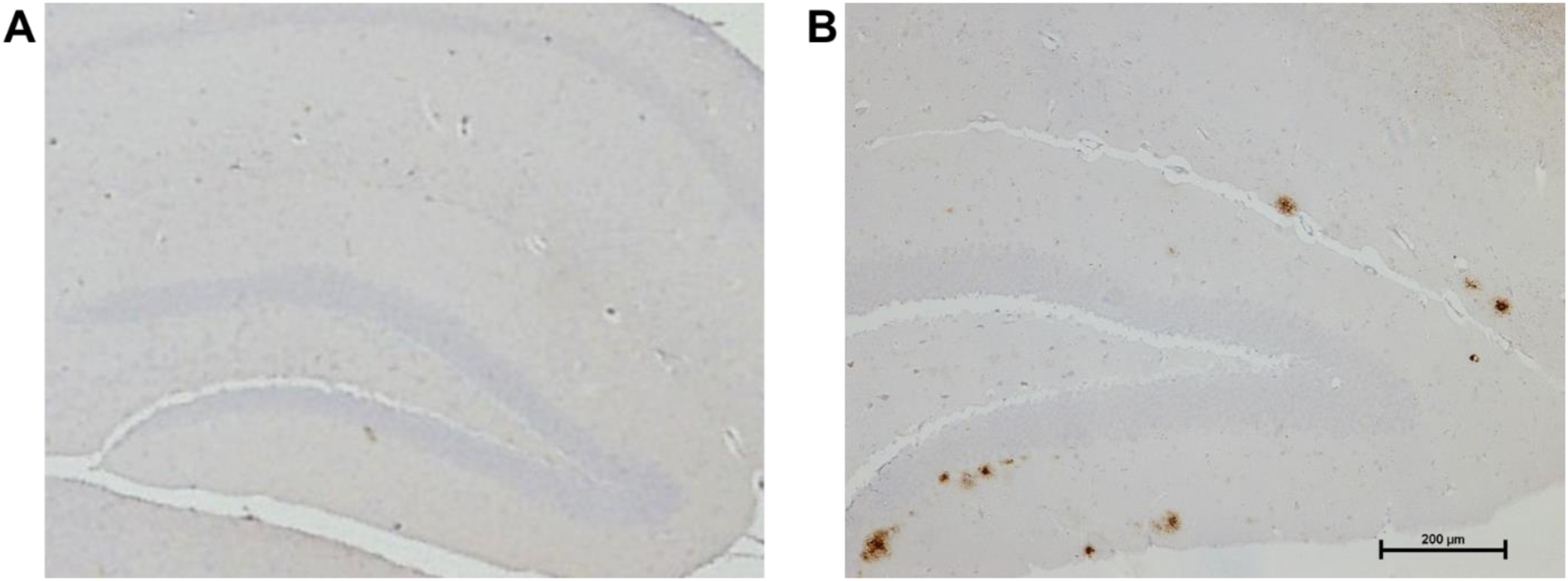
Representative hippocampal sections. A) WT mice do not develop any amyloid plaques. B) In the J20-AD mouse, diffuse plaques can be found predominantly forming in the dentate gyrus of the hippocampus by 9m, with one or two developing in the deep cortex, but not in the superficial layers.

## Discussion

The present study aimed to investigate neurovascular function in the J20-hAPP mouse model of AD to establish whether there were any haemodynamic and neural differences at early time points (6-9 months) compared to WT mice. Stimulation-evoked cerebral haemodynamics responses (blood volume and HbO) were significantly enhanced in J20-AD mice as measured in a stable chronic cranial window preparation (Figure 2). This consistent and remarkable finding was irrespective of stimulation duration (2s/16s) or gas condition (hyperoxia/normoxia). The washout of HbR was always smaller in J20-AD mice, but much smaller under normoxia. These responses were consistent across all experimental setups and the timing of experiments post-anaesthesia. These findings are contrary to the majority of the published literature with respect to neurovascular function in the J20-hAPP mouse^10–12^. However, one study has previously reported augmented haemodynamics in the APP/PS1 double transgenic model^13^. In that study, a multi-modal approach was used to investigate cerebral haemodynamics in response to electrical stimulation of the hind-paw. Laser-Doppler flowmetry, multi-photon laser scanning microscopy and intrinsic optical imaging (similar to 2D-OIS) all confirmed enhanced HbT in 7m APP/PS1 mice and not at earlier than 5m, similar to our data from the J20-AD mice between 6-9m of age. However, Kim and colleagues did not measure neural activity and speculated that the enhanced haemodynamic responses could be attributed to either increased neuronal activity or increased vascular reactivity. Our data along with others investigating neural activity confirm that there is neuronal hyperexcitability between 6-9m of age in the J20-hAPP mouse, but that there is no difference in vascular reactivity between WT controls and J20-AD mice as demonstrated by the response to hypercapnia. Previous research from our laboratory has shown no significant differences in either haemodynamics or neural activity between 9-12m in the same J20-mouse model unless an electrode is inserted into the brain^9^. Using exactly the same approach, we present here novel findings of hyperexcitability and enhanced evoked haemodynamic responses at an earlier age, between 6-9m, in the J20-mouse model of AD. Combined, these studies suggest that these effects are only seen at an earlier age point and are, therefore, time-dependent.

A major finding from this study was that under normobaric hyperoxia, the baseline blood volume and saturation across the entire cortex was substantially higher in J20-hAPP mice compared to control mice, and this effect disappeared upon transitioning to normoxia, where J20-AD mice displayed a significant decrease in HbT to match the baseline of WT controls (Figure 4). This enhanced blood volume could be seen across all vascular compartments acrossthe observable cortical regions. This oxygen-specific effect on cerebral haemodynamics was only observed in the J20-AD mice and was not a feature of WT mice and did not seem to affect stimulation-evoked haemodynamics in terms of HbT volume. In early stages of AD, soluble Aβ fragments can be cleared from the brain through paravascular pathways in the brain, including the glymphatic system^14^ and the intramural periarterial drainage system^15^. Naturally increasing cerebral blood flow may enhance the clearance of soluble Aβ in the early stages of AD onset. Since our data show that normobaric hyperoxia can elevate baseline blood volume across the cortex in young J20-AD mice, this mechanism may provide a potential disease prevention strategy enhancing the clearance of Aβ before plaques begin to form, as well as reducing the formation and concentration of Aβ oligomers. As we can visualise plaques from around the age the J20-hAPP mice, these observations could be related to early Aβ deposition. In terms of corroboration and translatability of these findings, it has previously been demonstrated that hyperbaric oxygen therapy reduces neuroinflammation in the 3xTg AD mouse model^16^. Here, we present findings that *normobaric* hyperoxia can enhance cerebral haemodynamics and therefore may potentially be able to clear Aβ from the brain.

Another finding from this study was that the J20-AD mice were hyperexcitable, as indicated by the enhanced MUA evoked by whisker stimulation (Figure 3) consistent with other studies showing neural hyperexcitability in the J20 mouse^17^. The cause of hyperexcitability in J20-AD mice, and indeed some AD patients, may be due to neurophysiological abnormalities caused by the presence of Aβ^18–21^, or by aberrant APP processing resulting in different intracellular domain fragments^22–24^. Unprovoked seizures can occur in 10-22% of AD patients^25^, especially myoclonic seizures, which are specifically due to the hyperexcitability of the cortex. For unknown reasons, the prevalence of seizures in AD-patients, including they myoclonic type, is more frequent in younger AD-patients, especially that of early-onset AD^26^. This hyperexcitability at a young age is also reflected in our data from the J20-AD mouse. Furthermore, another study showed a higher incidence of epileptiform-like discharges in the APP/PS1 mouse model compared to controls which were correlated to the number of Aβ plaques between 4-9m of age^27^. The hyperexcitability and epileptic activity of the J20-AD mice dissipates by the age of 8m^22^. Therefore, the hyperexcitability seen in the J20-AD mouse, APP/PS1 mice and younger AD-patients may be related to the brains initial reaction to the emerging presence of Aβ oligomers, or that of aberrant APP processing. Whisker deflection in APP/PS1 mice also resulted in much higher neural responses in the cortex, also supporting our findings^28^. In our study, the observed enhanced neural activity couples well with the enhanced haemodynamic responses, suggesting that neurovascular coupling in the J20-AD mouse is intact and functioning well at this younger age of 6-9m.

In our acute experimental sessions, the intactness of NVC was masked by the technical procedure used and a mismatch between neural activity and haemodynamic response was recorded (Figure S2). As described briefly^9^, insertion of an electrode into the mouse brain can cause cortical spreading depression (CSD) to occur across the cortex^29^. CSD is characterised by prolonged vasoconstriction that can persist for some time (in some cases over an hour) and this generally dampens haemodynamic responses and overall cerebral blood flow across a wide area of the cerebral cortex^30^. During this time, neural activity often recovers quickly, however the recovery rate of haemodynamics varies and there is a considerable lag^31^. As such, despite the enhanced MUA in J20-AD mice to stimulations, the haemodynamics measured are comparable to WT controls due to CSD. Although cerebral haemodynamics recover in both WT and J20-AD mice, the rate of recovery between these varies, and J20-AD mice take longer^9^. By the end of the experimental period, HbT increases in J20-hAPP mice to match that of WT controls (Figure S2). Our data highlight the major impact that acute imaging sessions have on neurovascular studies. Future experiments could combine 2D-OIS with non-invasive optical neural readouts (such as genetically encoded calcium indicators (GECIs) e.g. GCaMP in neurons, although these have their own technical and physiological considerations, discussed in depth elsewhere^32^.

A limitation of the study was that animals were lightly anaesthetised throughout. While it is increasingly argued that all neurovascular studies should be performed using awake animals^33^ due to the effects of anaesthesia on neurovascular function, previous research from our laboratory has determined an optimal anaesthetic regimen that has minimal effect on cerebrovascular function and reactivity^34^. Performing experiments at 1-hour post induction of anaesthesia, and maintaining sedation via low levels of isoflurane (<0.5%), is comparable to awake imaging in terms of haemodynamic responses and profiles to whisker stimulation^34^. Use of such an anaesthetic regime avoids the multiple confounds associated with behavioural state (arousal, locomotion, stress, grooming etc.), which are present in the awake animal. A second limitation is that there were no empirical values for tissue oxygenation both during baseline and during stimulations and therefore we assumed tissue O_2_ saturation to be at 70% and haemoglobin concentration to be 100µM. A third limitation was that blood flow measurements were not quantified and therefore we were unable to assess perfusion in the brain as a whole. Most other studies however indicate chronic hypoperfusion in patients and the J20-AD mouse^35^.

In conclusion, using a stable chronic imaging preparation, we have shown that, at early time points, the J20-hAPP mouse model of AD exhibits enhanced haemodynamics in the brain marked by an increased blood volume response to sensory stimulations compared to WT controls. The likely cause of such increased blood volume responses in young J20-AD mice is neural hyperexcitability, suggesting that neurovascular function is preserved in these mice. A key finding from this study was that under normobaric hyperoxia, baseline blood saturation and volume is enhanced in all vascular compartments in the brain of J20-AD mice. This effect did not present under normoxia, with transition from hyperoxia to normoxia resulting in a large decrease of HbT/blood volume in J20-AD, but not WT mice. The enhancement of blood volume in normobaric hyperoxia may provide a time-dependent therapeutic strategy by driving enhanced paravascular clearance pathways in the brain, potentially clearing soluble beta-amyloid and preventing the formation of plaques. Future work should investigate the effect of regular normobaric hyperoxia on the levels of beta-amyloid in the J20-AD mouse.

## Materials & Methods

### Animals

All animal procedures were performed with approval from the UK Home Office in accordance to the guidelines and regulations of the Animal (Scientific Procedures) Act 1986, as well as being approved by the University of Sheffield ethical review and licensing committee. 6-9m old male wild-type (WT) C57BL/6J mice (n=8) and male heterozygous transgenic J20-AD B6.Cg-Zbtb20Tg(PDGFB-APPSwInd)20Lms/2Mmjax) (MMRRC Stock No: 34836-JAX | J20) mice (n=9) were used. All mice were housed with littermates in a 12hr dark/light cycle at a temperature of 23C, with food and water supplied *ad-libitum*.

### Chronic Thinned Cranial Window Preparation

Mice were anaesthetised with 7ml/kg i.p. injection of fentanyl-fluanisone (Hypnorm, Vetapharm Ltd, UK), midazolam (Hypnovel, Roche Ltd, UK), diluted in sterile water in a 1:1:2 by volume ratio for surgery induction, and maintained in a surgical anaesthetic plane by inhalation of 0.5-0.8% isoflurane. Core body temperature was maintained at 36.5-37.0C through rectal temperature monitoring. Mice were placed in a stereotaxic frame (Kopf Instruments, US) and the scalp was excised. The bone overlying the right somatosensory cortex was thinned to translucency forming a thinned cranial optical window (measuring ∼9mm^2^). A thin layer of clear cyanoacrylate glue was applied over the cranial window to reduce specularities and to reinforce the window. Dental cement (Super Bond C&B, Sun Medical, Japan) was applied around the window to which a metal head-plate was chronically attached. All mice were given 2 weeks to recover before the first imaging session was performed.

### 2D-Optical Imaging Spectroscopy (2D-OIS)

2D-OIS measures changes in cortical haemodynamics by estimating changes in total haemoglobin (HbT), oxyhaemoglobin (HbO) and deoxyhaemoglobin (HbR) concentrations as described previously^36^. For the experimental session, mice were anaesthetised (as described above) and placed into a stereotaxic frame with heads fixed using attached headplates. Anaesthesia was maintained using low-levels of isoflurane (0.3-0.6%). For imaging, the right somatosensory cortex was illuminated using 4 different wavelengths of light appropriate to the absorption profiles of the differing haemoglobin states (495nm ± 31, 559nm ± 16, 575nm ± 14 & 587nm ± 9) using a Lambda DG-4 high-speed galvanometer (Sutter Instrument Company, US). A Dalsa 1M60 CCD camera with a frame rate of 32Hz operating in a 4×4 binning mode acquired images at 184×184 pixels; giving a pixel resolution of 75×75µm, was used to capture the re-emitted light from the cortical surface.

All spatial images recorded from the re-emitted light underwent spectral analysis based on the path length scaling algorithm (PLSA) as described previously^36,37^, which uses a modified Beer-Lambert law with a path-length correction factor converting detected attenuation from the re-emitted light with a predicted absorption value. Relative HbT, HbR and HbO concentration estimates were generated from baseline values in which the concentration of haemoglobin in the tissue was assumed to be 100µM and O_2_ saturation to be 70%. The spectral analysis produced 2D-images of micromolar changes in volume of HbT, HbO & HbR over each stimulation period.

### Stimulation Paradigm & Experimental Overview

A mechanical whisker stimulation paradigm was used. Whiskers were mechanically deflected for a 2s duration and a 16s duration at 5Hz using a plastic T-shaped stimulator which caused a 1cm deflection of the left-whisker pad in the rostro-caudal direction. Each individual experiment consisted of 30 stimulation trials (for 2s) or 15 stimulation trials (for 16s) from which a mean trial was generated after spectral analysis of 2D-OIS (as described previously). Stimulations were performed in 100% O_2_ (normobaric hyperoxia), a gas transition to medical air (normobaric normoxia; 21% O_2_) as well as an additional 10% CO_2_-hypercapnia test of vascular reactivity. The same set of experiments with the same timings were performed on all mice on all experimental days. 2-weeks post-surgery the 1^st^ imaging session was performed with 2D-OIS alone (a chronic session as imaging was through an intact skull). 1-week after the 1^st^ imaging session a 2^nd^ terminal imaging session was performed in combination with neural electrophysiology.

### Neural Electrophysiology

In order to assess neurovascular function in its entirety, both haemodynamic and neural measures were obtained. Simultaneous measures of neural activity alongside 2D-OIS were performed in a final acute imaging session 1-week after the 1^st^ imaging. A small burr-hole was drilled through the skull overlying the barrel cortex (as defined by the biggest HbT changes from 2D-OIS imaging) and a 16-channel microelectrode (100µm spacing, 1.5-2.7MΩ impedance, site area 177µm^2^) (NeuroNexus Technologies, USA) was inserted into the whisker barrel cortex to a depth of ∼1500µm. The microelectrode was connected to a TDT preamplifier and a TDT data acquisition device (Medusa BioAmp/RZ5, TDT, USA). All data collected was sampled at 24kHz and downsampled to 6kHz for analysis of multi-unit activity (MUA) and local-field potentials (LFPs).

### Region Analysis

Analysis was performed using MATLAB (MathWorks). An automated region of interest (ROI) was selected using the stimulation data from spatial maps generated using 2D-OIS. The threshold for a pixel to be included within the ROI was set at 1.5xSTD, therefore the automated ROI for each session per animal represents the area of the cortex with the largest haemodynamic response, as determined by the HbT. For each experiment, the response across all pixels within the ROI was averaged and used to generate a time-series of the haemodynamic response against time for HbT, HbO & HbR.

### Statistical Analysis

Statistical analyses were performed using GraphPad Prism v8. To compare haemodynamic & neural responses, statistical comparisons were made on HbT, HbO, HbR & MUA values using two-way repeated measures ANOVAs as well as 2-tailed unpaired t-tests. P-values <0.05 were considered statistically significant. All the data are presented as mean values ± standard error of mean (SEM), unless otherwise stated.

### Immunohistochemistry

At the end of terminal experiments, mice were euthanized with an overdose of pentobarbital (100mg/kg, Euthatal, Merial Animal Health Ltd) and transcardially perfused with 0.9% saline followed by 4% ice-cold paraformaldehyde (0.1M, pH7.4). Brains were dissected and embedded in paraffin wax. 5µm coronal sections were obtained using a cryostat. Immunohistochemistry was performed using an avidin-biotin complex (ABC) method (as described previously^5^). Briefly, sections were deparaffinised, rehydrated and quenched of endogenous peroxidase activity in 0.3% H_2_O_2_/methanol solution. Following antigen retrieval (pressure cooker at 20psi at 120C for 45s (pH6.5)) sections underwent additional pre-treatment in 70% formic acid. Sections were incubated with 1.5% normal serum followed by incubation with the primary antibody (biotinylated anti-Aβ – 1:100, BioLegend, USA) for 1 hour. Horseradish peroxidase avidin-biotin complex (Vectastain Elite Kit, Vector Laboratories, UK) was used to visualise antibody binding along with 3,3-diaminobenzidine tetrahydrochloride (DAB) (Vector Laboratories, UK). All sections were counterstained with haematoxylin, dehydrated and mounted in DPX. Sections were imaged using a Nikon Eclipse Ni-U microscope attached to a Nikon DS-Ri1 camera.

## Acknowledgments

We would like to thank Prof Lennart Mucke (Gladstone Institute of Neurological Disease & Department of Neurology, UCSF, CA, US) as well as the J. David Gladstone Institutes for the J20-hAPP mice. We wold like to thank Michael Port for building and maintaining the whisker stimulation device and 2D-OIS apparatus.

## Supplemental Data

**Figure S1.**
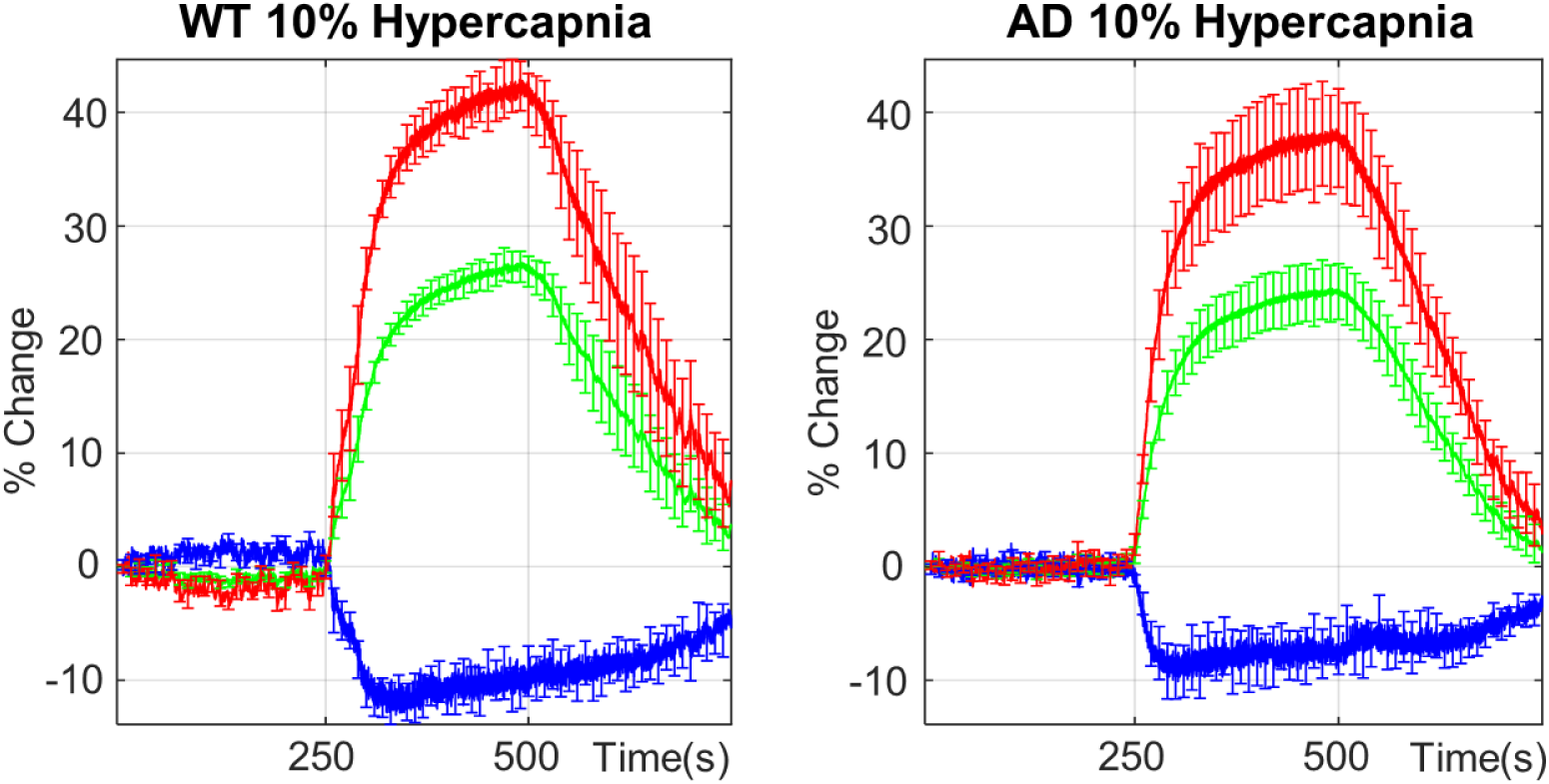
10% Hypercapnia Responses. Chronic data for WT (n=8) and J20-AD (n=9) hypercapnia responses showed no significant differences between groups. Hypercapnia was performed in 100% oxygen.

**Figure S2.**
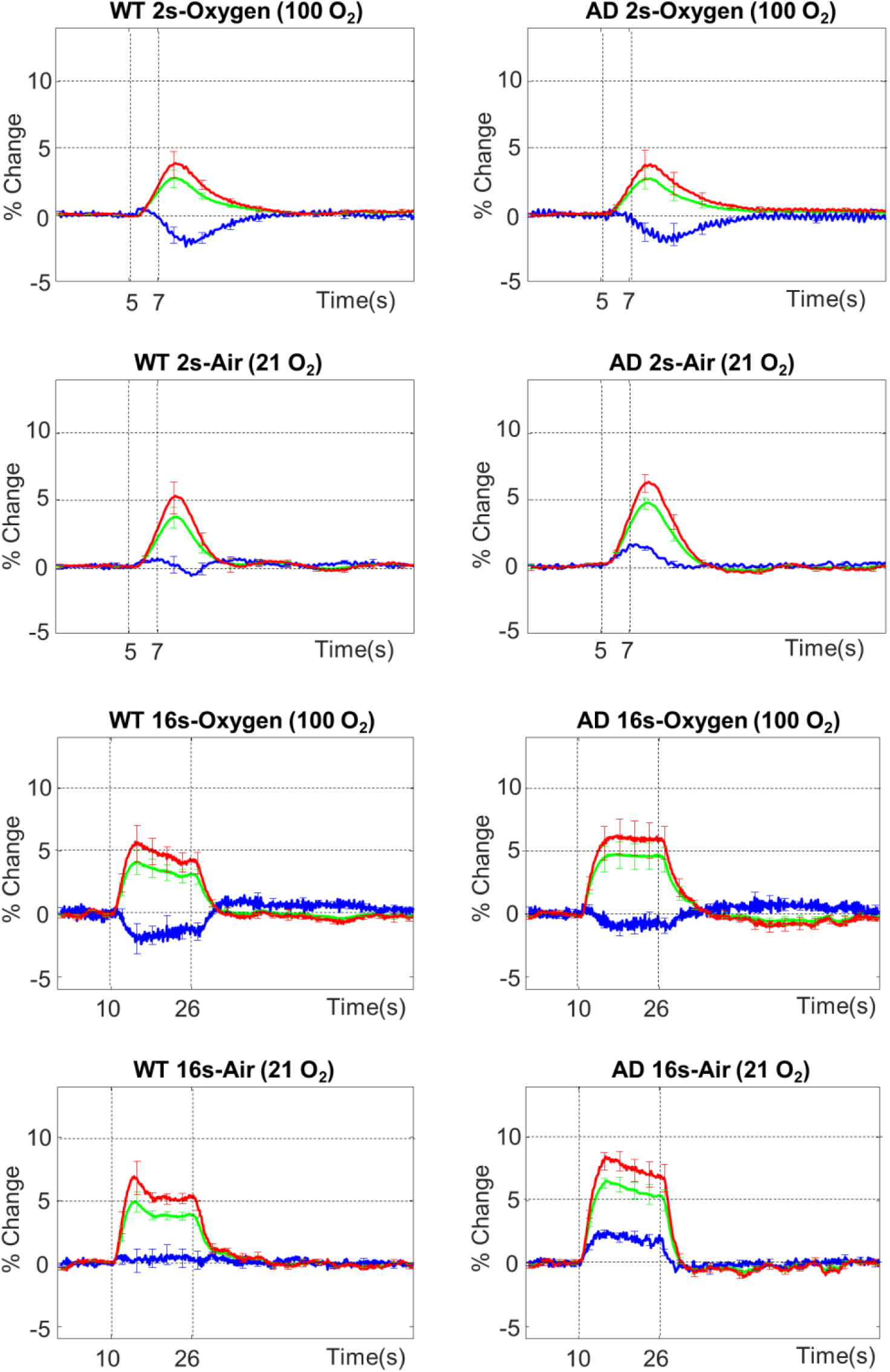
Haemodynamic Responses During Acute Imaging Session (with Electrode Inserted). There are no significant differences in HbT between WT (n=6) and J20-AD (n=5). HbT/blood volume recovers in both groups as a result of time post-electrode insertion. Washout of HbR is impaired under 21% oxygen conditions in both WT and J20-AD mice.

